# Integrating Enviromics to Predict Performance and Guide Clonal Deployment in Eucalyptus spp.

**DOI:** 10.64898/2025.12.01.691191

**Authors:** João Gabriel Zanon Paludeto, Gustavo Eduardo Marcatti, Regiane Abjaud Estopa, Jaroslav Klápště, João Carlos Bespalhok-Filho, Rafael Tassinari Resende

## Abstract

Genotype by environment interaction (G×E) remains a central challenge for tree breeding, as it is difficult to extrapolate trial results to untested sites and complicates confident genotype deployment. Enviromics, by integrating environmental covariates into predictive models, offers a way to overcome these limitations and guide clonal deployment. This study evaluated an enviromic framework applied to 15 Eucalyptus spp. clones across 5,189 inventory plots located in southern Brazil. 10,000 Engineered Enviromic Markers (EEMs) were built from 3,869 soil, climate, and remote sensing covariates, using random forest and integrated into a mixed-model ensemble to predict mean annual increment standardized at seven years (MAI7) across the Target Population of Environments (TPE). Predictive accuracy was assessed through a Leave-One-Region-Out cross-validation procedure. The enviromic model achieved a higher performance than a baseline G×E model, with Pearson and Spearman correlations above 0.90 and a root mean squared error (RMSE) of 3.09, compared to 0.45–0.39 correlations and RMSE of 8.73 for the baseline. Spatial predictions enabled the delineation of breeding zones that minimized G×E, while also revealing regions with high discriminant power for testing new genotypes. We also applied a two-step clonal deployment procedure combining enviromic predictions with a frost-risk penalization map, refining recommendations for frost-prone areas. When comparing recommended versus planted clones in inventory plots, the framework indicated an average expected productivity gain of 13.4%. These results demonstrate the potential of enviromics as a decision-support tool for clonal deployment, enhancing productivity while accounting for environmental risks, and paving the way for future multi-omics integration.

## 1 INTRODUCTION

Genotype by environment interaction (G×E) has always been a major concern for forest tree breeders (Costa e Silva et al., 2006; Hardner et al., 2010; Rosado et al., 2012; Santos et al., 2016). The G×E can often pose significant challenges in plant breeding, as it influences the estimates of genetic gain, reducing overall heritability, and impacting selection accuracy of the affected traits (Li et al., 2017). Combined with the current speed of climate change, G×E represent a continuous challenge for properly defining the Target Population of Environments (TPE) (Cappa et al., 2015; Cooper et al., 2021; Ray et al., 2022; Cruz et al., 2025). Moreover, its magnitude also plays a major role in deployment strategies. When G×E is strong, it prevents reliable extrapolation of trial results to untested sites, which in turn complicates the spatial allocation of genetic material across heterogeneous environments (Crossa et al., 2021; Marcatti et al., 2017; Resende et al., 2025, 2021).

Throughout the history, several analytical tools have been developed to quantify, interpret and analyze G×E in crops. Early approaches were based on linear regression of genotype means on environmental indices (Finlay and Wilkinson, 1963) and on the sum of square decomposition to derive stability parameters (Eberhart and Russell, 1966; Shukla, 1972), later some new approaches emerged, notably the Additive Main Effects and Multiplicative Interaction (AMMI) model (Gauch, 1992) and the Genotype plus Genotype by Environment (GGE) biplot (Yan et al., 2000). With advancements of quantitative genetics and the popularization of mixed effect models, BLUP-based indices were introduced to simultaneously evaluate stability and productivity of genotypes (Resende, 2004). More recent approaches have also contributed to refining G×E analysis, such as the Weighted Average of Absolute Scores (WAASB), which combines AMMI with BLUPs, bringing G×E investigation to a mixed-model framework (Olivoto et al., 2019). Still, regardless of whether based on ANOVA or mixed models, these approaches remain restricted to interpret and understand G×E for a specific set of tested environments, since they do not incorporate explicit soil and climate covariates, limiting their ability to predict performance on untested conditions (Costa-Neto et al., 2022).

The possibility to predict performance and, as a result, allocate genotypes optimally to untested sites has been a subject of immense interest in both plant and animal breeding (Resende et al., 2021). To achieve this goal, Heslot et al. (2014) and Jarquín et al. (2014) introduced the concept of genome-based reaction-norms model, which uses several Environmental Covariates (ECs) to model G×E in a genomic prediction framework. Since then, subsequent studies have demonstrated that these approaches can contribute to a higher prediction accuracy of genotype performance on untested sites (Pérez-Rodríguez et al., 2015; Cuevas et al., 2016; Acosta-Pech et al., 2017; Marcatti et al., 2017; He et al., 2019). With the increasing numbers of ECs to characterize the environment, this approach quickly became what we know these days by enviromics (Xu, 2016).

The classical Enviromics approach is using reaction-norm kernel models, proposed by Jarquín et al. (2014), a method very similar to the classic GBLUP (VanRaden, 2008) where, instead of a realized genomic relationship matrix between individuals, a linear kernel of environmental relationships (covariances) is built from the ECs, linking phenotypic variation to specific environmental factors to model genotype-specific response curves over continuous environmental gradient. The model provides the ability to predict genotype performance in untested locations or years, thereby supporting and optimizing genotype deployment (Costa-Neto et al., 2022; Resende et al., 2025).

In GIS-based spatial prediction approaches (GIS: Geographic Information System), proposed by Marcatti et al., (2017) and then refined to an enviromics context by Resende et al. (2021), any defined land area is a grid of georeferenced pixels compiled into GIS raster layers, so that each grid cell (pixel) has an associated environmental profile built from the ECs. The initial step is the development of Engineered Enviromic Markers (EEMs) associating ECs with a particular trait using machine learning algorithms. The EEMs are then used as predictor variables to fit reaction norm mixed models and predict for the whole TPE. By modelling G×E interactions at a granular pixel level, this approach provides spatial predictions that support both breeding decisions (e.g., definition of breeding zones, where G×E is minimized) and deployment strategies, exploring G×E and, consequently enhancing realized genetic gain through a pixel-wise optimized deployment of genotypes (Araújo et al., 2024; Bahia et al., 2025; Cruz et al., 2025; Marcatti et al., 2017; Resende et al., 2025).

Although analytical methods exist to incorporate ECs into G×E studies, very few works have directly applied these models in operational breeding programs using real tree inventory data and focusing on genotype deployment (Marcatti et al., 2017; Scolforo et al., 2020, 2017). Furthermore, there are still no consolidated pipelines capable of delivering pixel-wise genotypic recommendations across continuous areas.

Given this context, the objective of this study is to evaluate an integrated GIS-based approach using ECs and mixed model framework, combined with real *Eucalyptus* spp. inventory data, to predict clonal performance across the defined Target Population of Environments (TPE), delivering a decision-support tool capable of optimize clonal allocation.

## 2 METHODS

### 2.1 Study area and phenotypic data

The study analyzed 15 distinct *Eucalyptus* spp. clones from five species (*E. urophylla* x *E. grandis*, *E. dunnii*, *E. saligna*, *E. benthamii* and *E. urophylla*) (Figure 1b) across 5,189 inventory plots from the Klabin company, distributed in different areas of the Brazilian states of Paraná (PR), São Paulo (SP), and Santa Catarina (SC). The focus was on assessing the mean annual increment (MAI7) productivity variable, which measures volume per hectare per year (m^3^ ha^−1^ yr^−1^) standardized to a seven-year growth period. Each clone had a specific and well-fitted equation to project volume per hectare from the current age to age 7 and consequently obtained MAI7.

**Figure 1 –.**
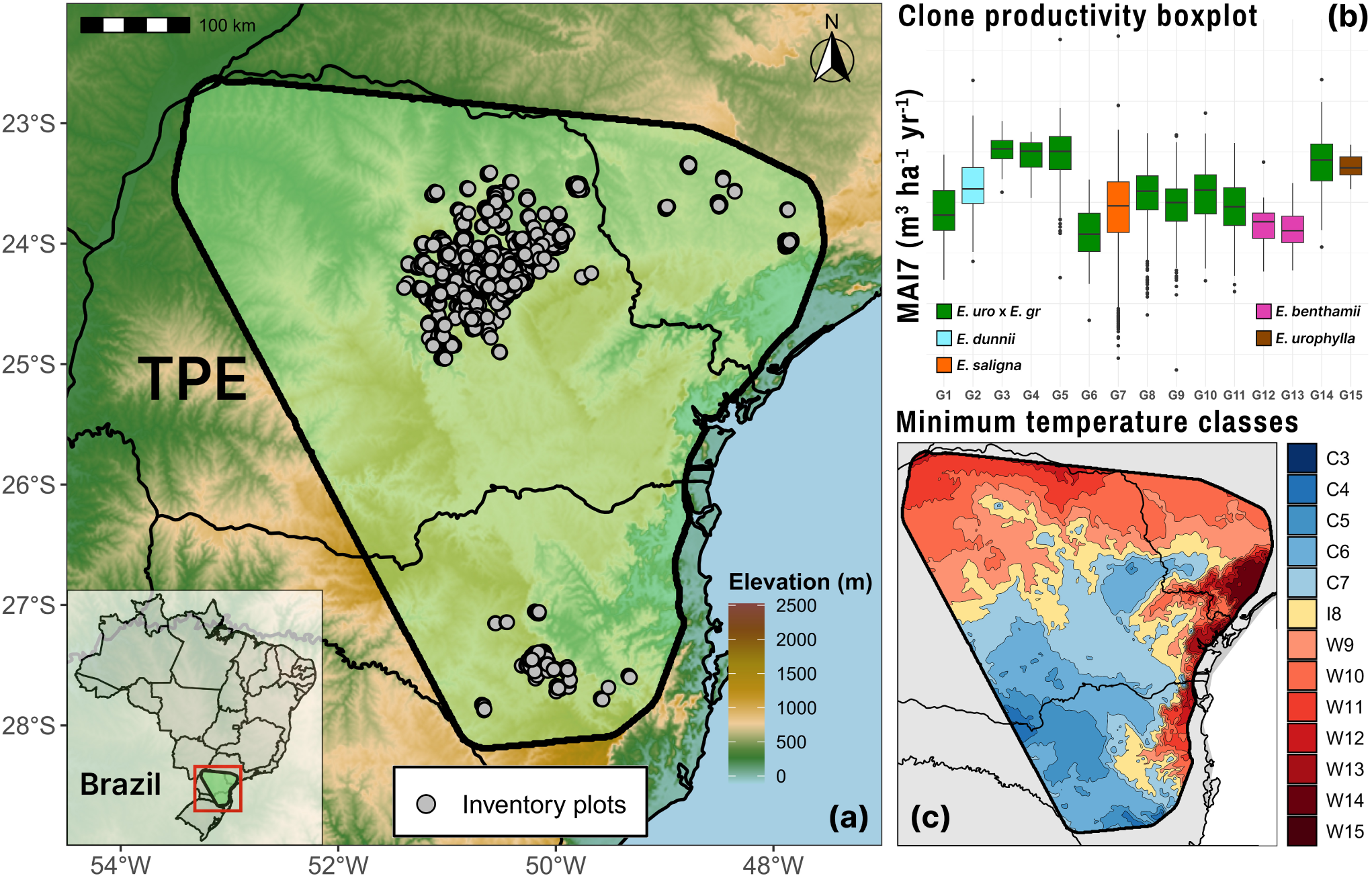
Study area located in south Brazil **(a)** TPE encompassing the states of Paraná (PR), São Paulo (SP), and Santa Catarina (SC), with 5,189 inventory plots. **(b)** Boxplot of productivity (MAI7 m^3^ ha^−1^ yr^−1^) of each evaluated clone for five *Eucalyptus* species (hybrids of *Eucalyptus urophylla* x *E. grandis*, and the pure species *E. dunnii*, *E. saligna*, *E. benthamii* and *E. urophylla*); **(c)** Zoning of the TPE based on minimum temperature classes used to guide clone recommendations according to frost susceptibility. Classes C3–C7 (blue tones) represent colder regions with higher frost frequency and severity, where lower class numbers correspond to lower minimum temperatures. Class I8 (yellow) indicates intermediate conditions with moderate frost risk. Classes W9–W15 (red tones) represent warmer regions, where higher numbers correspond to higher minimum temperatures and negligible frost risk.

**Figure 2 –.**
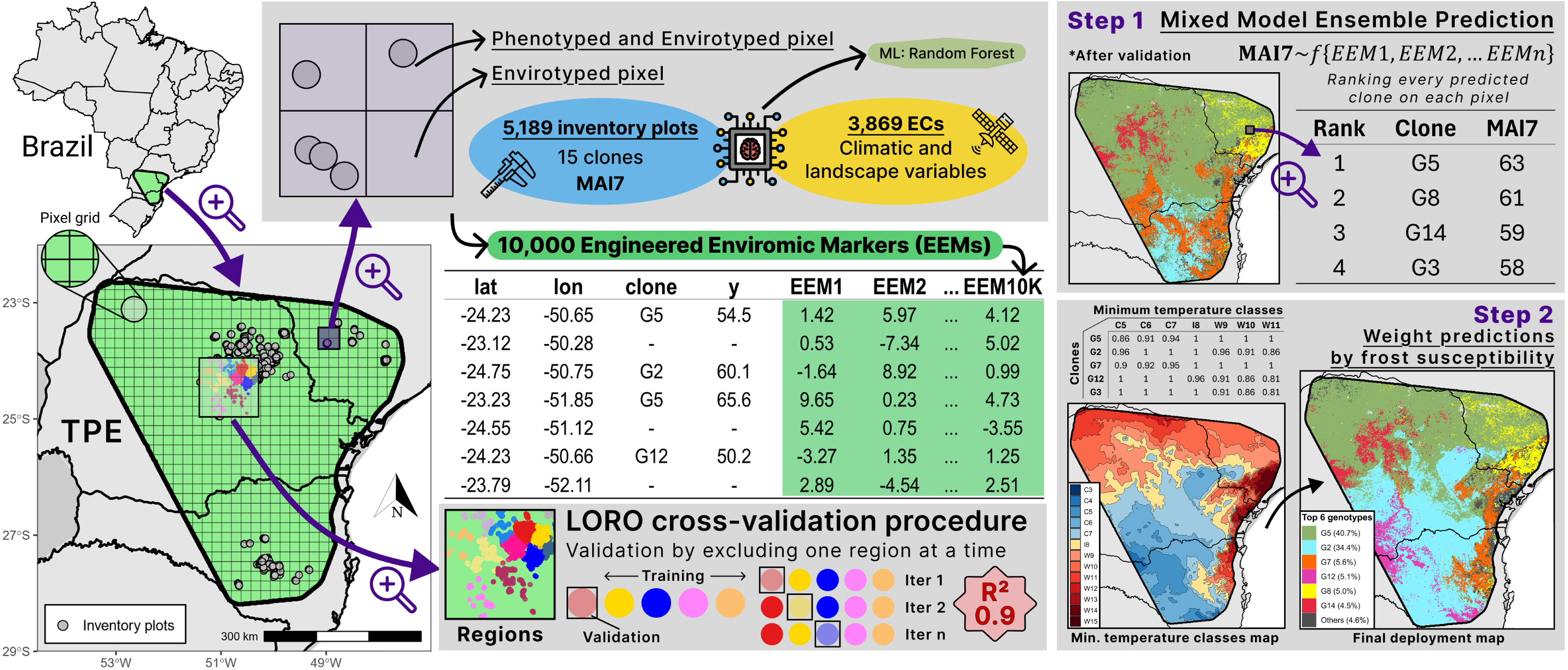
Detailed application of the enviromics methodology, from data phenotyping and envirotyping to a final two-step prediction considering weighted predictions by frost susceptibility.

The Target Population of Environments (TPE) was delineated according to Cruz et al. (2025), by connecting the outermost forest inventory plots, generating a geospatial polygon that defines the core area of interest. A 50 km buffer was then applied beyond this boundary to account for surrounding conditions and extrapolation potential (Figure 1a). The prediction grid was constructed using approximately 1 km² spatial resolution (1 × 1 km pixels), resulting in 208,589 individual pixel bins. All spatial analyses were conducted using the WGS84 coordinate reference system, in alignment with international geospatial standards.

### 2.2 Envirotyping and Engineered Enviromic Markers (EEM)

Environmental data for the study were envirotyped from five primary sources, resulting in a total of 3,869 ECs which were spatially downloaded and aggregated over the Target Population of Environments (TPE).

**a) MODIS Satellite:** Contributed with 2,070 covariates. This satellite-based platform provides detailed images and data on vegetation cover and surface temperature, essential for understanding factors like photosynthesis and transpiration. Data were collected for tile h13v11 from January 1, 2016, to December 31, 2023, using the pyModis library (pypi.org/project/pyModis).
**b) NASA POWER:** Supplied 26 covariates. This data source offers information on solar radiation and air temperature, which are essential for assessing energy input and evapotranspiration rates in crops. Data was collected from 1990 to 2022 and 2006 to 2022, using the requests library (https://pypi.org/project/requests).
**c) SoilGrids:** Contributed with 244 covariates, including soil density, cation exchange capacity, and organic carbon content. This data provides insights into soil fertility and structure, which are critical for plant growth. The data covers depths up to 200 cm and includes mean values as well as 0.05, 0.5, and 0.95 percentiles. Data was accessed using the SoilGrids library (https://pypi.org/project/soilgrids).
**d) WorldClim:** Provided 20 bioclimatic variables, such as monthly temperature and rainfall. This platform is crucial for capturing annual climate trends, seasonality, elevation and extreme weather conditions, which can significantly impact crop performance. Data was downloaded also using the requested library, referenced above.
**e) ERA5:** Supplied 1509 covariates. This high-resolution reanalysis product provides hourly climate data, including precipitation, temperature, soil moisture, wind speed, and radiation fluxes. These variables are essential for capturing short- and long-term environmental dynamics that affect plant growth. Data were retrieved from 2009 to 2021 using the Climate Data Store API (cdsapi library, https://pypi.org/project/cdsapi).

Following the methodology proposed by Resende et al. (2025), the Engineered Enviromic Markers (EEM) were developed through a Random Forest (RF) model using Python’s scikit-learn library (pypi.org/project/scikit-learn). The RF model, consisting of *N* regression trees (function *T*), employs the residual sum of squares to make predictions (*m*) based on environmental covariates (*W*). Each tree is constructed using a random subset of data (*i* ∈ *I*) with specific parameters (*j* ∈ *J*), enhancing the model’s overall robustness. This procedure generated 10,000 predictors, with environmental means (*m*) calculated from randomly chosen genotypes to reflect the influence of environmental variability on genetic performance. Finally, the predictors were grouped into eight distinct clusters using hierarchical clustering, where the n^th^ regression tree is fitted as:

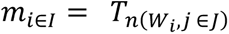

Where the *k^th^* EEM (*∈_k_*) was obtained by aggregating similar predictors using RF, and defined as:

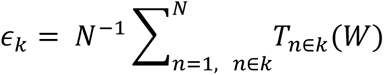

Two strategies were used to validate the EEMs: the first involved assessing the correlation between observed and predicted values, while the second employed leave-one-out cross-validation to evaluate the model’s predictive ability for unsampled environmental pixels. In the leave-one-out approach, the model is trained using data from all inventory plots except one, which is then predicted. This process is repeated for each plot in the dataset.

### 2.3 Predictive modelling and cross-validation

To predict genotype performance within the TPE, we implemented an ensemble-based random regression framework integrating EEMs predictors. For each ensemble iteration, a subset of these EEMs was randomly sampled, one per cluster, and used to fit a mixed-effects model as random slopes for each genotype:

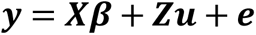

where ***y*** is the phenotypic observations vector; ***X*** is the incidence matrix for fixed effects (including the intercept); ***β*** is the vector of fixed effects; ***Z*** is the incidence matrix of random effects; ***u*** is the vector of random genetic terms (intercepts and reaction norms for each EEM) and 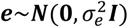.

The random effects ***u***~***N***(**0**, **Σ ⊗ *I***) follow a multivariate normal distribution, where **Σ** is the variance-covariance matrix of the genetic terms across EEMs. For each genotype, the reaction norm is modeled by a random intercept and a set of random slopes corresponding to the selected EEMs. Each iteration (totaling 100 runs) produced genotype-specific coefficients, which were applied to a grid of 1 km² resolution across the defined Target Population of Environments (TPE). This grid consisted of 208,589 spatial pixels, and predictions were generated for all genotypes at each location. The ensemble predictions were then averaged across runs to obtain stable estimates of MAI7.

To assess predictive accuracy, a leave-one-region-out (LORO) cross-validation strategy was applied. The dataset was clustered into 20 spatial colored regions based on geographic proximity (Figure 3a). At each fold, one region was excluded from model training and used for validation.

**Figure 3 -.**
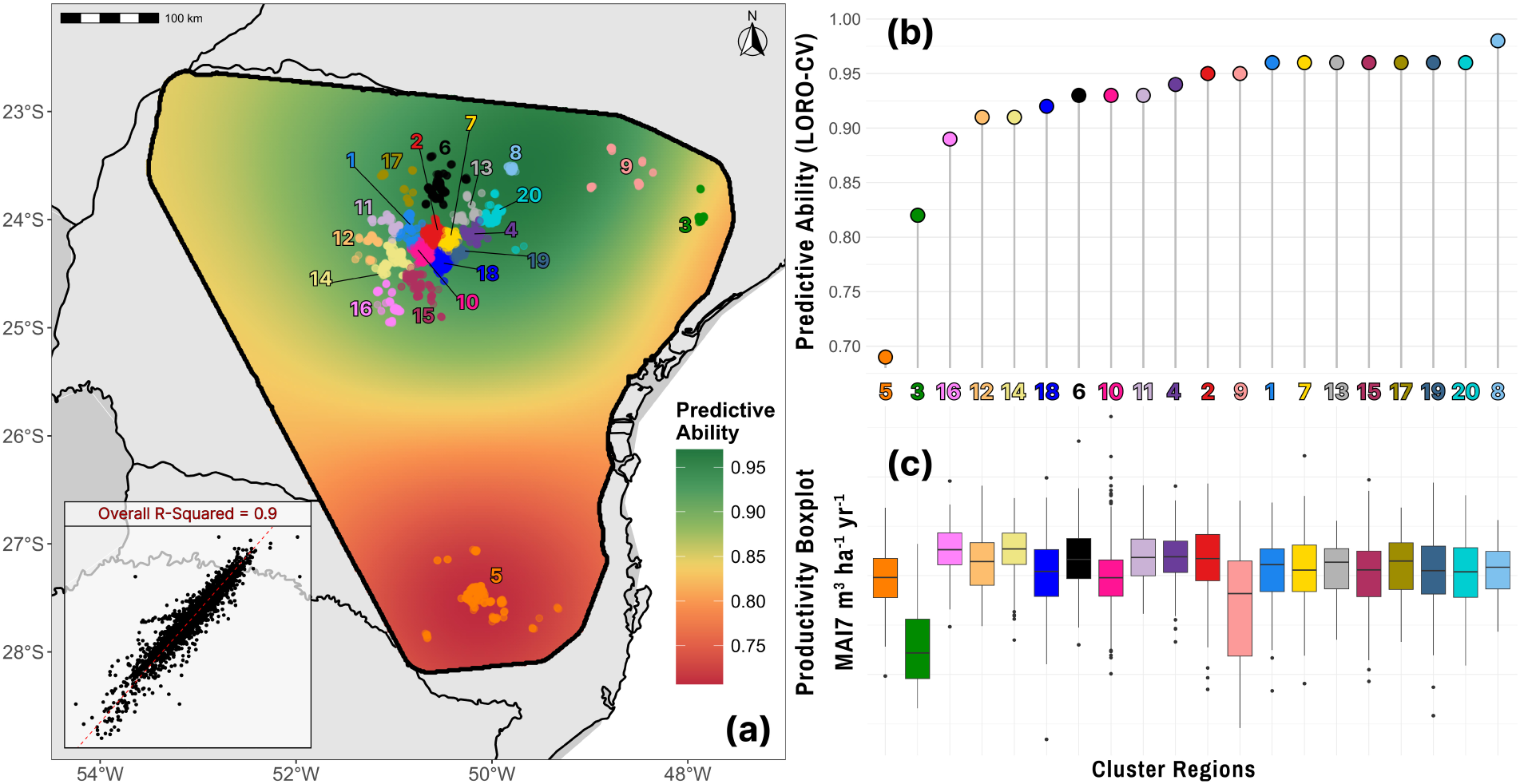
Spatial and region-level evaluation of the enviromic model performance. **(a)** Predictive ability interpolated across the TPE using ordinary kriging, cluster-colored regions defined by geographical proximity and used in the Leave-One-Region-Out (LORO) cross-validation procedure. **(b)** Predictive ability obtained for each region. **(c)** Boxplots of mean annual increment at 7 years (MAI7, m^3^ ha^−1^ yr^−1^) per region.

To assess whether the model was effectively capturing genotype-by-environment responses, rather than predominantly reflecting environmental effects, a “false genotype prediction test” was applied. For each region validation fold, each pixel bin was predicted using the reaction norm of a random genotype that was not present in that pixel. The predicted value was then compared against the observed phenotypic value of the actual genotype located at that specific bin.

To establish a reference for the enviromic model performance, we fitted a baseline G×E model with only the intercept as fixed effect. Random terms include region, genotype, and genotype by cluster interaction. No ECs or EEMs were used in this model. All model evaluation metrics based on LORO cross-validation, whether for the baseline model, the enviromics method or the false genotype prediction test, were assessed using squared Pearson and Spearman correlation coefficients, along with the Root Mean Squared Error (RMSE).

To evaluate the spatial performance of the enviromic model, we applied ordinary kriging to interpolate the squared LORO predictive ability across the entire TPE. First, the mean predictive ability of each cluster was associated with its geographic centroid. A regular prediction grid was then created over the study area, and a theoretical variogram was fitted to the squared correlations using a Gaussian model. The interpolated surface was plotted as a continuous heatmap representing an expected local prediction accuracy.

### 2.4 Two-step clonal deployment

To determine the most suitable clone for each pixel within the Target Population of Environments (TPE), we implemented a two-step recommendation procedure. First, enviromic predictions of clone performance were obtained. Then, predicted values were adjusted using a penalization factor derived from a temperature-based zoning map, which guides the deployment of genetic materials according to their frost susceptibility (Figure 1c).

To estimate the productivity gain associated with the two-step clonal deployment recommended by the enviromic framework, we compared, at pixels that contained inventory plots, the predicted performance of the recommended clone with the observed productivity of the clone that was actually planted.

### 2.5 Breeding Zones

To identify Breeding Zones (BZs), defined as regions within the TPE that minimize G×E (Resende et al., 2021), the prediction grid was aggregated from 1 km² to 10 km² resolution to facilitate interpretation and alignment with operational deployment scales, and, for each bin, mean predicted MAI7 values were calculated for each genotype. Then, according to (Resende et al., 2025) we computed a distance matrix from a pairwise correlation matrix among pixels and submitted to hierarchical clustering using the Ward method. The TPE was then partitioned into four breeding zones, following the suggestion of a silhouette metric, each maximizing the similarity within zones.

### 2.6 Patterns of predicted yield

To investigate the spatial patterns of both environmental potential and genetic variability across the TPE, two complementary maps were derived based on the set of predicted values for all genotypes. First, the environmental potential of each location was estimated by averaging the predicted productivity (MAI7) across all EEMs. Second, the genetic variance for each location was computed based on the variance EEMs. Also, to evaluate the G×E patterns across the predicted TPE, we constructed reaction norm curves for each clone.

## 3 RESULTS

Model performance was evaluated using a Leave-One-Region-Out (LORO) cross-validation strategy, in which the inventory plots were subdivided into 20regions by geographic distance, that served as independent validation units. With validation strategy the baseline model yielded overall predictive abilities of 0.45 and 0.39 for the Pearson and Spearman correlation coefficients, respectively, with a Root Mean Squared Error (RMSE) of 8.73. As the baseline model was fitted only once, standard deviations were not available for any of its metrics. The false genotype prediction procedure resulted in higher predictive ability than the baseline method, with mean Pearson and Spearman correlations of 0.57 ± 0.23, and a lower RMSE of 7.04 ± 1.50. The enviromics model presented the highest overall performance, with Pearson and Spearman correlation coefficients of 0.92 ± 0.02 and an RMSE of 3.09 ± 0.32.

Across regions, the false genotype prediction scenario showed higher standard deviations and correlation coefficients ranging from 0.43 to 0.68. In contrast, the predictive ability of the enviromic model remained consistently high, with Pearson correlations ranging from 0.69 to 0.98 and low associated standard deviations.

For a macro view of the predictive abilities considering the spatial variation for the entire TPE, Figure 3a was generated using the Kriging procedure, which was supported by the cluster predictive abilities for the enviromic model, as presented in Table 1 and Figure 3b.

**Table 1 -.** Predictive abilities estimated, using Pearson and Spearman correlations for each cluster region using the Leave-One-Region-Out cross validation procedure for the evaluated models.

| Region | Baseline* |  |  | Enviromics |  |  | False Genotype Prediction |  |  |
| --- | --- | --- | --- | --- | --- | --- | --- | --- | --- |
|  | Pearson | Spearman | RMSE | Pearson | Spearman | RMSE | Pearson | Spearman | RMSE |
| 1 | 0.59 ± NA | 0.53 ± NA | 7.08 ± NA | 0.96 ± 0.01 | 0.95 ± 0.01 | 2.44 ± 0.23 | 0.65 ± 0.16 | 0.61 ± 0.19 | 6.62 ± 1.45 |
| 2 | 0.64 ± NA | 0.60 ± NA | 7.53 ± NA | 0.95 ± 0.01 | 0.96 ± 0.01 | 2.78 ± 0.32 | 0.56 ± 0.22 | 0.52 ± 0.24 | 7.40 ± 1.31 |
| 3 | 0.19 ± NA | 0.21 ± NA | 20.18 ± NA | 0.82 ± 0.01 | 0.84 ± 0.01 | 7.23 ± 0.35 | 0.55 ± 0.28 | 0.59 ± 0.26 | 12.00 ± 3.26 |
| 4 | 0.51 ± NA | 0.47 ± NA | 7.06 ± NA | 0.94 ± 0.01 | 0.93 ± 0.02 | 2.27 ± 0.24 | 0.52 ± 0.19 | 0.51 ± 0.19 | 5.86 ± 0.87 |
| 5 | 0.45 ± NA | 0.45 ± NA | 12.36 ± NA | 0.69 ± 0.14 | 0.70 ± 0.16 | 6.24 ± 2.29 | 0.57 ± 0.37 | 0.54 ± 0.38 | 6.58 ± 2.27 |
| 6 | 0.35 ± NA | 0.21 ± NA | 8.20 ± NA | 0.93 ± 0.01 | 0.92 ± 0.02 | 2.52 ± 0.18 | 0.59 ± 0.22 | 0.57 ± 0.22 | 5.94 ± 0.99 |
| 7 | 0.62 ± NA | 0.55 ± NA | 7.05 ± NA | 0.96 ± 0.01 | 0.96 ± 0.01 | 2.62 ± 0.15 | 0.60 ± 0.17 | 0.60 ± 0.17 | 7.37 ± 1.42 |
| 8 | 0.35 ± NA | 0.22 ± NA | 6.02 ± NA | 0.98 ± 0.01 | 0.98 ± 0.01 | 1.49 ± 0.16 | 0.56 ± 0.24 | 0.55 ± 0.22 | 5.43 ± 1.08 |
| 9 | 0.38 ± NA | 0.49 ± NA | 15.04 ± NA | 0.95 ± 0.02 | 0.94 ± 0.02 | 4.35 ± 0.65 | 0.68 ± 0.25 | 0.67 ± 0.26 | 10.20 ± 2.71 |
| 10 | 0.60 ± NA | 0.43 ± NA | 6.77 ± NA | 0.93 ± 0.01 | 0.93 ± 0.02 | 3.00 ± 0.20 | 0.57 ± 0.15 | 0.55 ± 0.16 | 7.11 ± 1.02 |
| 11 | 0.65 ± NA | 0.53 ± NA | 6.07 ± NA | 0.93 ± 0.01 | 0.93 ± 0.01 | 2.56 ± 0.19 | 0.45 ± 0.36 | 0.45 ± 0.32 | 6.32 ± 1.59 |
| 12 | 0.22 ± NA | 0.06 ± NA | 8.36 ± NA | 0.91 ± 0.01 | 0.89 ± 0.01 | 3.12 ± 0.16 | 0.59 ± 0.21 | 0.52 ± 0.24 | 6.22 ± 1.14 |
| 13 | 0.46 ± NA | 0.42 ± NA | 6.32 ± NA | 0.96 ± 0.02 | 0.95 ± 0.01 | 1.95 ± 0.33 | 0.51 ± 0.28 | 0.51 ± 0.30 | 6.00 ± 1.36 |
| 14 | 0.56 ± NA | 0.46 ± NA | 7.47 ± NA | 0.91 ± 0.01 | 0.91 ± 0.01 | 2.80 ± 0.13 | 0.51 ± 0.19 | 0.49 ± 0.20 | 5.99 ± 1.06 |
| 15 | 0.39 ± NA | 0.33 ± NA | 8.83 ± NA | 0.96 ± 0.01 | 0.96 ± 0.01 | 2.81 ± 0.20 | 0.62 ± 0.21 | 0.63 ± 0.21 | 7.88 ± 1.81 |
| 16 | 0.27 ± NA | 0.22 ± NA | 9.51 ± NA | 0.89 ± 0.01 | 0.90 ± 0.01 | 3.12 ± 0.11 | 0.43 ± 0.24 | 0.44 ± 0.24 | 6.29 ± 0.93 |
| 17 | 0.08 ± NA | 0.16 ± NA | 8.38 ± NA | 0.96 ± 0.01 | 0.97 ± 0.01 | 1.99 ± 0.15 | 0.65 ± 0.21 | 0.68 ± 0.19 | 5.57 ± 1.28 |
| 18 | 0.58 ± NA | 0.54 ± NA | 7.66 ± NA | 0.92 ± 0.01 | 0.93 ± 0.01 | 3.55 ± 0.17 | 0.54 ± 0.19 | 0.53 ± 0.20 | 8.17 ± 1.52 |
| 19 | 0.60 ± NA | 0.49 ± NA | 7.77 ± NA | 0.96 ± 0.01 | 0.96 ± 0.01 | 2.68 ± 0.17 | 0.67 ± 0.15 | 0.67 ± 0.14 | 7.33 ± 1.36 |
| 20 | 0.57 ± NA | 0.52 ± NA | 6.98 ± NA | 0.96 ± 0.01 | 0.97 ± 0.01 | 2.29 ± 0.07 | 0.67 ± 0.26 | 0.67 ± 0.26 | 6.62 ± 1.60 |
| Overall | **0.45 ± NA** | **0.39 ± NA** | **8.73 ± NA** | **0.92 ± 0.02** | **0.92 ± 0.02** | **3.09 ± 0.32** | **0.57 ± 0.23** | **0.57 ± 0.23** | **7.04 ± 1.50** |

\*The baseline model produces a single prediction per pixel and, therefore, does not have a standard deviation associated.

Figure 4a illustrates the performance curves, i.e. reaction norms for all genotypes, highlighting the top five performing clones with distinct colors. These genotypes exhibited similar response patterns. As expected, all clones increse their overall predicted productivity along the environmental gradient. Notably, clone G5 displayed the highest overall productivity and a significantly higher productivity in the worst environments.

**Figure 4 -.**
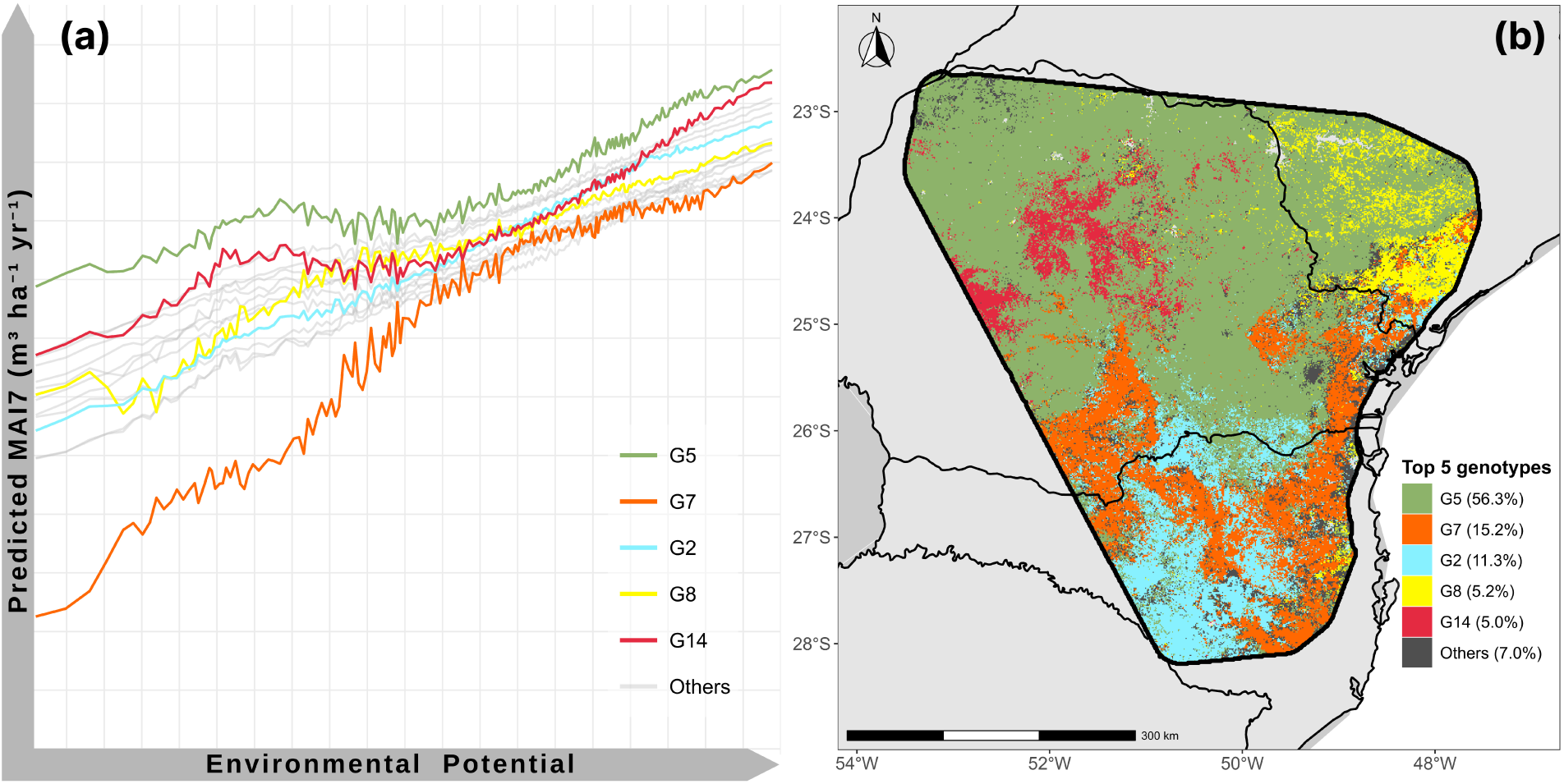
G×E specific prediction across the environmental gradient and spatial prediction of best-performing clones in the Target Population of Environments (TPE). **(a)** Reaction norm curves for the top 6 recommended genotypes based on predicted MAI7 values (m^3^ ha^−1^ yr^−1^) across an environmental potential gradient. (**b**) Spatial distribution of the top-performing clone per pixel in the TPE.

The spatial deployment considering only the best-performing genotype per pixel within the TPE according to the enviromic prediction can be visualized in Figure 4b. The five top recommended clones (G5, G7, G2, G8 and G14) accounted for 93% of the area, with G5 dominating in the PR state portion of the predicted area and accounting for 56.3% of the entire TPE.

The two-step clonal deployment, which combines enviromic predictions (Figure 4b; Figure 5a) with frost risk penalization (Figure 2c; Figure 5b), produced a refined deployment map (Figure 5d). The recommended area for G2 increased from 11.3% to 34.4%, while G5 decreased from 56.3% to 40.7% and G7 from 15.2% to 5.6%. Also, G12 emerged as a new recommended genotype, covering 5.1% of the TPE. The comparison between the predicted performance of the recommended clones on the pixels that contain inventory plots versus the productivity of the clones currently planted indicated an overall expected gain of 13.4% in mean annual increment (MAI7) (Figure 5c).

**Figure 5 -.**
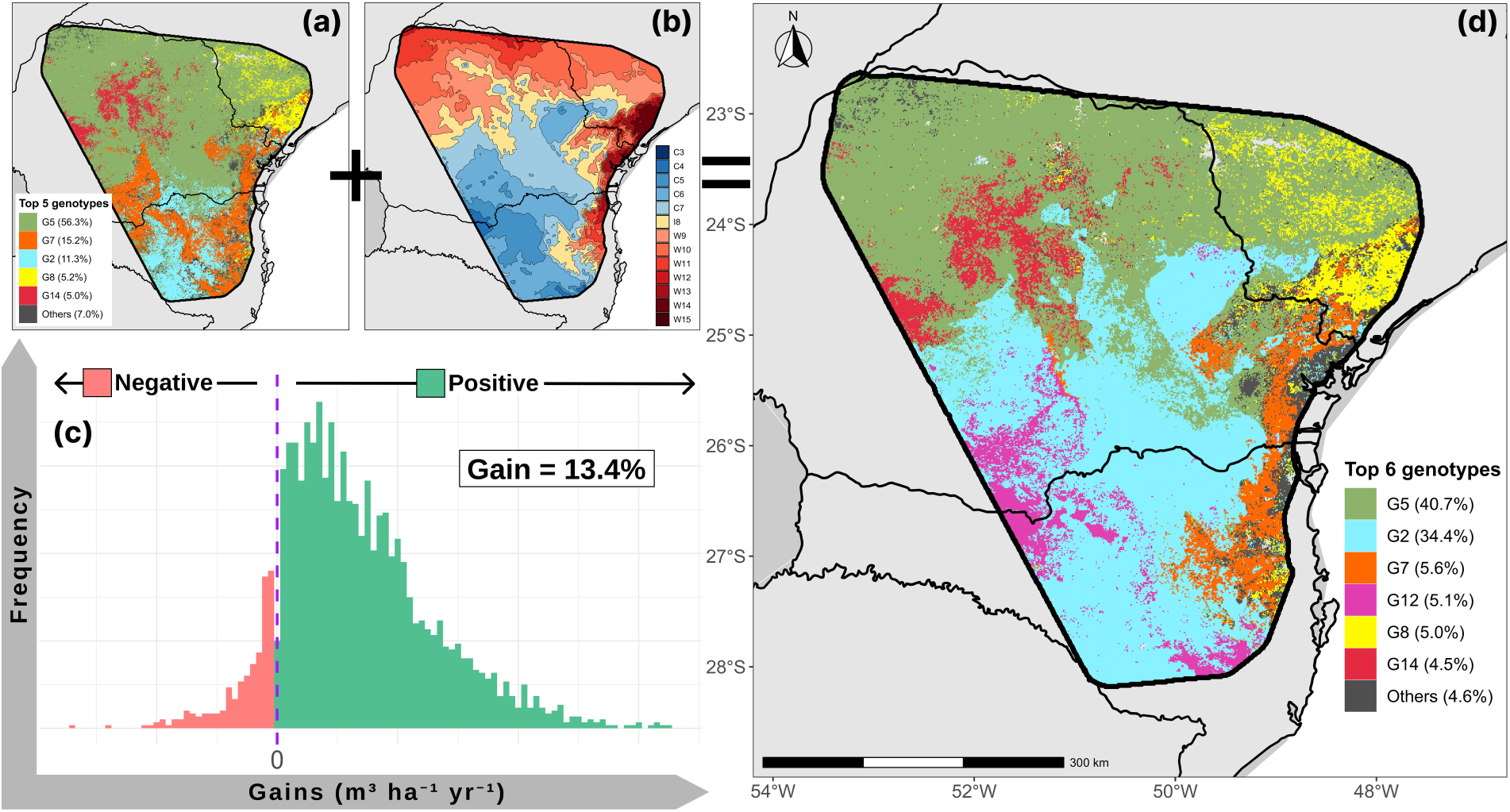
Two-step recommendation procedure for clonal deployment across the Target Population of Environments (TPE). **(a)** Enviromic predictions of clonal performance, indicating the top-performing genotypes per pixel. **(b)** Minimum temperature-based zoning map defining frost risk classes. ***(c)*** Histogram of positive and negative gains when comparing the predicted performance of the recommended clone with the productivity of the planted clone on inventory plots. **(d)** Final deployment map where predicted performance was adjusted by penalization factors according to frost susceptibility.

The yield potential for MAI7 and genetic variances of predicted yield for each clone are in Figure 6. A clear spatial pattern was observed, with the highest environmental potential concentrated in the central region of the TPE, especially in areas surrounding the majority of inventory plots (Figure 1a; Figure 3a). In contrast, regions in the northwestern and northeastern extremes, exhibited not only the lowest environmental potential (Figure 6a), but also the highest genetic differentiation among predicted clones (Figure 6b).

**Figure 6 -.**
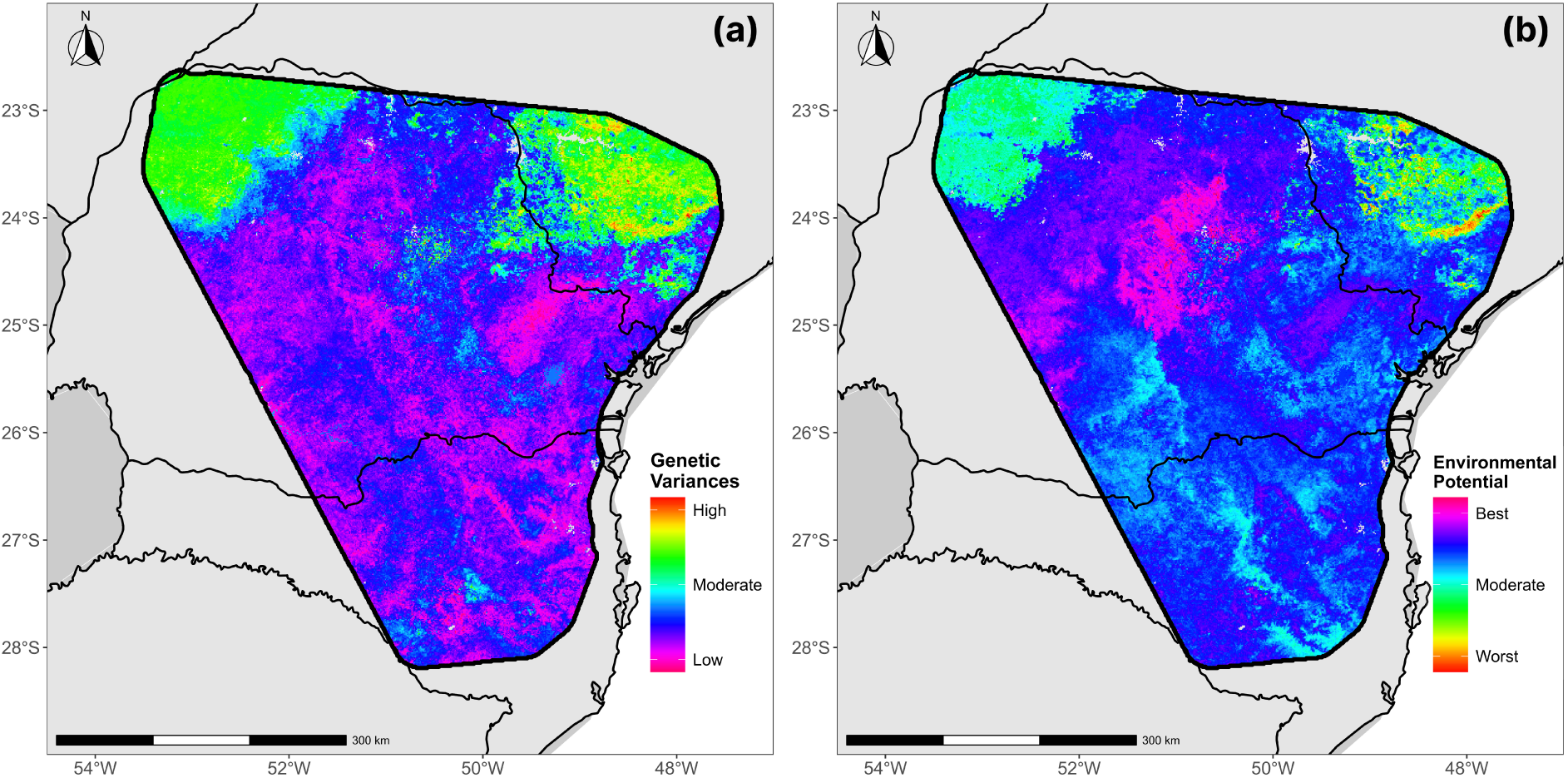
Environmental potential map **(a)** and genetic variances map **(b)** across the Target Population of Environments (TPE).

The resulting breeding zones (BZ1 to BZ4) are displayed in Figure 7, revealing a clear spatial structure across the landscape. BZ1 and BZ2 covered the northern and central portions of the TPE, respectively, while BZ3 and BZ4 encompassed more environmentally contrasting southern and coastal areas.

**Figure 7 –.**
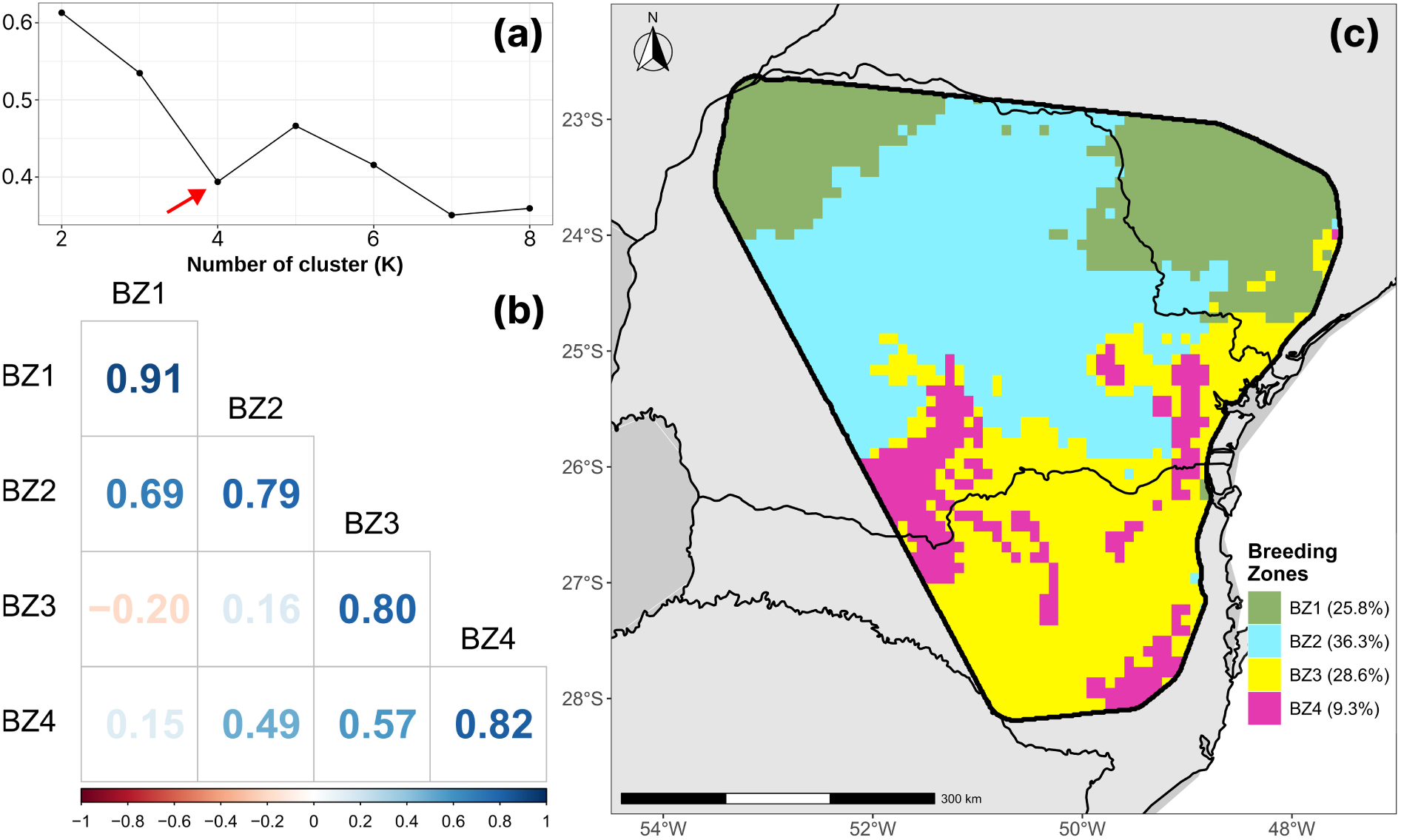
Breeding zones within the Target Population of Environments (TPE). **(a)** Silhouette metric used to define the optimal number of breeding zones. **(b)** Correlation matrix within and between each defined breeding zone. **(c)** Spatial distribution of the four breeding zones assigned to 10 km-resolution bins across the TPE.

The genetic correlations between and within zones are summarized in Figure 7b. Breeding zones presented high within correlations, ranging from 0.79 to 0.91 for BZ2 and BZ1, respectively. Between zone correlations varied markedly, with some pairs exhibiting moderate values (BZ1–BZ2: 0.69; BZ3–BZ4: 0.57), while others presented low (BZ2–BZ3: 0.16; BZ1–BZ4: 0.15) or even negative correlations (BZ1–BZ3: −0.20).

## 4 DISCUSSION

### 4.1 From Environment to Interaction: Insights from Baseline and Enviromic Models

The comparison of modeling strategies clearly highlights the value of incorporating environmental information in prediction of tree growth across sites. The baseline model yielded moderate predictive power, indicating that traditional G×E modeling, as expected, can predict performance, at some point. However, it was clearly outperformed by the enviromic model that integrated explicit envirotypic covariates with genotype data. The enviromic approach achieved notably higher prediction accuracy with consistently low standard deviations, suggesting that, the use of environmental descriptors, such as the Engineered Enviromic Markers (EEMs) used in this study enable the model to explain genotype performance differences across each site with robust and stable predictions across clusters (Costa-Neto et al., 2022; Jarquín et al., 2017; Resende et al., 2025; Xu et al., 2022). By modeling reaction norms of genotypes over continuous environmental gradients, enviromic models better capture G×E interactions and can even predict performance in new future environments (Figure 4), something that the baseline method is not capable of (Callister et al., 2024; Resende et al., 2025).

Clusters 3 and 5 showed the poorest predictive performance among all regions, with the lowest predictive abilities and highest RMSE values in both the baseline and enviromics models (Figure 3). For the baseline, predictive abilities were approximately 0.20 for region 3 and 0.45 for region 5, with RMSE values of 20.18 and 12.36, respectively. The same happened for the enviromics model, which, although still achieving high predictive abilities, these regions exhibited the lowest predictive abilities (~0.83 and ~0.69) and comparatively higher RMSE values (7.23 ± 0.35 for cluster 3 and 6.24 ± 2.29 for cluster 5). Cluster 3 is located northeastern extreme of the TPE, while cluster 5 lies farther south. They are both geographically distant from the core region where most training data were concentrated (Figure 1a; Figure 3a). This spatial disjunction resulted in a reduced representation of local G×E patterns during model training, disturbing the prediction accuracy in these areas, since, the more similar is an untested environment to the tested environment, the higher the chances of a good and assertive prediction (Araújo et al., 2024). This distance from the core region, and consequently, lower predictive abilities for these two clusters, are what dictate the patterns presented in the kriging procedure in Figure 3. Although the values are still high, they can be improved with more data on those regions, or even specific enviromic models for each of these areas.

The “false genotype prediction” scenario provides additional insight into the role of environment in prediction. In this approach, the reaction norm of a random genotype, not present in a specific pixel bin, was used to predict performance and compare to the genotype that is present in that bin. As expected, this mis-specified model performed substantially worse than the properly integrated enviromic model and, interestingly, better than the baseline model, in both predictive ability and RMSE. This actually reveals some important insight about the enviromic prediction framework. In a way, this test essentially predicts environmental performance alone, ignoring the actual G×E interaction. The improvement over the baseline method only strength the enviromic method and confirms that it is being successful in capturing environmental signals.

Moreover, the overall predictive ability of the false genotype prediction test was 38% lower than that of the enviromic model. In other words, this test serves as indirect evidence that the enviromic model is indeed capturing genotype-specific responses and not merely environmental patterns. Notably, regions 3 and 5, which performed poorly in both baseline and enviromic predictions, showed similar predictive ability in the false genotype test when compared to other clusters. This further supports the idea that the environmental component is being accurately modeled by the Engineered Enviromic Markers (EEMs).

This study relies on a small number of genotypes from several *Eucalyptus* species, so the observed genotype preferences across the environmental landscape may partly reflect species-level differences rather than within-species variation. In contrast, applying enviromics within an operational, single-species forest tree breeding programme introduces additional complexities. Unlike typical proof-of-concept studies that assess only a few genotypes, a nucleus-style breeding scheme maintains a broad, non-clonal base population to preserve diversity, alongside a cloned elite tier that may include hundreds of genotypes differing only subtly (Dungey et al., 2009; Wu et al., 2016). Furthermore, because each genotype is typically evaluated in only a subset of field trials, genetic connectedness between sites is often uneven, complicating the reliable estimation of genotype-by-environment interaction (Callister et al., 2024; Li et al., 2018). Genomic information can partly strengthen across-trial connectivity and improve G×E inference, but it does not fully overcome this limitation (Callister et al., 2021).

As a result, the predicted enviromic response surface for the elite genotypes may be fragmented, with individuals showing localized performance peaks and troughs across the deployment landscape. If these predicted peaks are used as the primary deployment criterion, the resulting strategy may be overly sensitive to uncertainties in environmental mapping layers. Given that operational deployment decisions must remain effective over decades, it may be more strategic to prioritise stability of performance across environmental gradients rather than maximising point performance at narrow optima (McKeand et al., 1990; Pupin et al., 2018). In this sense, using enviromics for “robustness screening” rather than for “peak chasing” has the potential to deliver more reliable gains under real-world deployment constraints.

### 4.2 Enviromics as a strategic tool in clonal deployment under frost-risk conditions

The heterogeneity of environmental conditions that impact genotype performance is a constant challenge to plant breeders of any species (Cooper and Messina, 2021; Ray et al., 2022; Scolforo et al., 2020). Understanding genotype by environment interaction (G×E) is pivotal for effective plant breeding, as it enables a precise deployment of genotypes across the different environmental condition, maximizing performance and better matching genotypes to specific environments (Cooper and Messina, 2021; Jarquín et al., 2017; Resende et al., 2025). In this scenario, incorporating climatic covariates into clonal deployment, matching the most suitable genetic material with the correct site will always offers considerable productivity gains (Callister et al., 2024; Marcatti et al., 2017; Scolforo et al., 2017). These gains can be further enhanced through the implementation of powerful tools like enviromics, which enable data-driven deployment strategies at granular resolutions, as demonstrated in this study, where predictions were made at an approximately 1km resolution. When integrated into deployment pipelines or optimization algorithms, such models can act as strong decision-support systems. In fact, in the present study, when comparing the predicted performance of the most suitable genotype for each site, i.e., replacing the currently planted clone in each pixel with the one indicated by the enviromic model, we estimated an average gain of 13.4% in operational productivity (Figure 5c).

The clonal deployment according only to the enviromic prediction across the TPE (Figure 4b) reveals key insights into the expected genotype adaptation patterns and the model’s capacity to capture G×E variation. The predominance of G5 in nearly half of the TPE (56.3%) emphasizes its broad adaptability and competitive advantage, particularly under environments with low potential (Figure 4a). Every top-performing clone closely followed the environmental potential gradient. However, substantial re-ranking was observed along the gradient, evidencing the presence of genotype-by-environment interaction (G×E) patterns. Clone G5 exhibited overall superiority throughout the whole gradient, particularly, suggesting a high degree of plasticity and robustness. Still, it is essential to emphasize that the environmental gradient represented in the x-axis corresponds to the average predicted MAI7 of each pixel using only EEMs, which was a proxy for environmental quality. Therefore, pixels from distant geographic regions (e.g., southern and northern extremes) may share similar average productivity values but differ in specific environmental characteristics, which obviously affects the relative performance of genotypes. Consequently, the superiority of G5 along the average gradient in Figure 4 does not necessarily imply universal adaptability to all regions with similar environmental potential.

However, it is important to emphasize that recommendations provided by the enviromic model are not absolute. Some biotic and abiotic risks, such as frost events or localized pest and disease outbreaks, are difficult to quantify accurately through statistical models alone. Thus, the expertise of the breeder and the strategic insight of the operational planning team will always remain indispensable. Their role is to interpret model outputs with caution, incorporating empirical knowledge with regional context to ensure robust and assertive deployment decisions. In other words, the key to maximize productivity is the synergy between data-driven decisions and the expert judgement of good professionals. With that in mind, we further refined enviromic predictions by considering clonal susceptibility to frost using a minimum temperature classes map in a two-step enviromic approach, since frost events are common in southern Brazil and can negatively affect productivity of susceptible genotypes.

As expected, a redistribution on clonal deployment could be observed when applying the two-step approach. G2, an *E. dunnii* clone, a species tolerant to mild frosts, was favored in moderately cold zones (C7–C5) displacing G5 (*E. urophylla × E. grandis*) and G7 (*E. saligna*), which, despite surviving milder-frosts, can often experience reduced productivity when exposed to these events. Another important subject is the emergence of an *E. benthamii* clone (G12) in high frost-prone areas (C5–C3). In these areas, even the tolerant G2 begins to lose performance, creating space for G12 under harsher frost conditions, as *E. benthamii* is widely known for its high frost tolerance (Oberschelp et al., 2022). In contrast, intermediate (I8) and warm zones (W9–W15) remained essentially unchanged, since the previously recommended clones (i.e., G5, G8, G14; all *E. urophylla × E. grandis* clones) are already adapted to more tropical and warm conditions. It is noteworthy that the penalization weights applied in each minimum temperature class were not assigned solely based on general species trends, as there is significant variability in frost tolerance within each species. Instead, they reflect clone-specific knowledge derived from operational experience, which is routinely incorporated into the company’s deployment decisions.

In practical deployment, the penalization process should also account for planting windows across different periods of the year. Seasonal variation in frost occurrence means that the same site can present very different levels of risk depending on the planting date. Thus, integrating temporal information into the penalization framework enhances the robustness of recommendations, aligning spatial allocation with long-term climatic patterns and the whole seasonal dynamics that influence ramets early survival and subsequent productivity.

Further refinements of the prediction framework could also incorporate additional biotic and abiotic risk factors. For example, the occurrence of pests and diseases may strongly affect genotype performance and could be mapped similarly to frost susceptibility (Alvares et al., 2017; Wingfield et al., 2008). Another interesting possibility is the integration of climate forecasts, including large-scale phenomena such as El Niño and La Niña, which could allow anticipating drought or excess rainfall events that not only affect but also alter the incidence of pests and diseases (Nóia Júnior et al., 2019). Incorporating these sources of risk into enviromic predictions would provide a more comprehensive decision-support tool, enabling not only the allocation of genotypes according to average productivity potential and G×E but also ensuring their resilience and productivity under future stress scenarios.

### 4.3 Discriminant Power and Breeding Zone Delimitation

It is extremely important to test new genotypes in regions that have a high discriminant power. In other words, areas that allow genotypes to fully express their potential, and consequently, induce high genetic variability, which is essential for identifying top-performing genotypes (Prus and Piepho, 2021). Although the concept of discriminant power has been discussed indirectly since the 1930s (Yates and Cochran, 1938), it was Yan et al. (2007) who formalized and popularized the term when proposing GGE Biplot and AMMI models to interpret G×E. Nowadays, enviromics can extend this concept to the whole TPE, by enabling genotype yield predictions at a pixel level. This approach not only reveals regions with high discriminant power among tested sites, but also identifies new breeding frontiers, where selection of superior genotypes may be more effective.

As illustrated in Figure 6b, the northern and northeastern extremes of the TPE exhibited higher genetic variance among predicted clones, suggesting a greater discriminant power in these areas. Interestingly, these regions were the ones that presented the lowest average potential productivity (Figure 6a). This result corroborates what was reported by Resende et al. (2024), who, when applying enviromics to maize, found that areas with lower-yield potential showed greater genetic variances than areas with high-yield potential. This highlights the importance of considering those areas on the detection of superior genotypes.

Another important outcome of enviromic prediction is the definition of breeding zones (BZs) in a TPE level (Costa-Neto and Fritsche-Neto, 2021; Resende et al., 2021). The main objective of subdividing the TPE into mega-environments (i.e., breeding zones) is to identify repeatable G×E patterns and reduce interaction within each zone. Although this approach increases the complexity of breeding programs, it enables more precise genotype targeting and can increase trait heritability within zones. As a result, it facilitates more accurate and effective selection, accelerates genetic gain, and improves both the efficiency of selection and the strategic deployment of trials (Callister et al., 2024; Gauch and Zobel, 1997; Heslot et al., 2014; Windhausen et al., 2012).

The within-zone correlations, which ranged from 0.79 to 0.91, indicate that the clustering method was effective in grouping environmentally and genetically similar regions, resulting in breeding zones with high internal homogeneity. This finding is consistent with Callister et al. (2024), who successfully classified their environments and obtained within-zone correlations ranging from 0.76 to 0.84. In contrast, BZ3 and BZ4, while moderately correlated with each other (0.57), showed low or even negative correlations with the other breeding zones (e.g., −0.20 between BZ1 and BZ3), reflecting a complex G×E structure across the TPE. These zones are located predominantly in the southernmost regions, characterized by cooler temperatures and higher latitudes. It is also noteworthy that the regions with low potential productivity and high discriminant power (i.e. genetic variance), as discussed earlier in this same section (Figure 6), were precisely those delineated as BZ1 in the clustering procedure (Figure 7c).

When G×E is significant, selecting genotypes within more homogeneous subregions can outperform broader selection strategies. Moreover, these environmentally distinct zones open opportunities for customized strategies, including the development of regionally adapted genotypes (Bustos-Korts et al., 2022). In other words, rather than being a limitation, a proper delimitation of breeding zones can open new opportunities by enabling breeding programs to diversify their genotype portfolio, target specific traits and deploy trials more effectively across the TPE.

Given the modeling framework applied in this study, one can argue that the identified zones may be more accurately described as *deployment zones* rather than strictly *breeding zones*. Unlike zones defined from multi-environment trials, which are designed to guide breeding selection, the present clustering was based solely on inventory data. As such, these zones are particularly suited to guide deployment strategies, supporting decisions on where to allocate specific clones across the TPE. Nevertheless, they still capture underlying G×E structures and can, therefore, serve as a complementary reference for breeding, while their primary utility remains in operational deployment.

It is also important to recognize that breeding zones are not static concepts, especially under a climate change scenario. Long-term climate variability and land-use can modify environmental gradients and change G×E patterns (Ukrainetz et al., 2018). Therefore, continuous re-evaluation of breeding zones, along with periodic refitting of enviromic models, becomes essential. This process not only ensures that recommendations remain aligned with current environmental realities but also enables breeding programs to detect climate trends and anticipate scenarios by focusing selection efforts on the development of genotypes adapted to emerging conditions (Callister et al., 2024; Cooper et al., 2023).

## 5 CONCLUDING REMARKS AND FUTURE PERSPECTIVES

From a broader perspective, the evolution of enviromic approaches in G×E modeling is evident. Modern reaction-norm frameworks that incorporate envirotypic data effectively move G×E analysis from mere interpretation to predictive application, allowing breeders to anticipate genotype performance in untested locations. This proactive use also expands the potential for guiding genotype deployment strategies, providing a powerful framework to match genotypes with specific climatic or management preferences to a more assertive breeding or deployment region. Furthermore, the granular spatial predictions enabled by enviromic models and powered by GIS methods, capture G×E patterns that would be missed by region-based classifications.

The proposed framework can still be further refined and extended beyond operational deployment. One promising direction is its integration with breeding trial data. In a bivariate context, combining data from inventory plots with breeding trials would enable more targeted analyses, allowing predictions of clonal behavior of breeding trials in inventory conditions. Such an approach could strengthen inference about G×E while leverage the interpretation of experimental tests (Resende, 2025).

Beyond this, enviromics also offers natural synergies with other omics approaches. Genomic and phenomic data provide complementary information on genetic architecture and phenotypes, which, when combined with environmental descriptors, can deliver a truly multi-omics predictive framework. Such integration could accelerate the development of robust genotypes adapted to both current and future environmental scenarios.

This study demonstrates the value of an enviromic-based approach for clonal recommendation in forestry. By combining high-resolution genotype performance prediction with a frost-risk penalization step, we refined deployment recommendations across a heterogeneous Target Population of Environments (TPE). Our approach helps bridge the gap between quantitative G×E prediction and on-the-ground management, delivering a decision-support tool that optimizes clonal allocation, boosts productivity, and leverages environmental heterogeneity at a commercial scale. By improving both the precision of immediate deployment decisions and the understanding of G×E, enviromic-based clonal recommendation can reshape tree breeding and deployment strategies, fostering more productive and climate-resilient forestry operations for the future.

## CRediT authorship contribution statement

**João Gabriel Zanon Paludeto:** Conceptualization, Data Curation, Formal analysis, Investigation, Methodology, Software, Visualization, Writing – original draft, Writing – review & editing; **Gustavo Eduardo Marcatti:** Conceptualization, Investigation, Methodology, Resources, Supervision, Software, Writing – review & editing; **Regiane Abjaud Estopa:** Conceptualization, Investigation, Validation, Writing – review & editing; **Jaroslav Klápště:** Investigation, Writing – review & editing; **João Carlos Bespalhok-Filho:** Supervision, Validation, Writing – review & editing; **Rafael Tassinari Resende:** Conceptualization, Investigation, Methodology, Project administration, Resources, Supervision, Software, Writing – review & editing.

